# The impact of ROI extraction method for MEG connectivity estimation: practical recommendations for the study of resting state data

**DOI:** 10.1101/2023.06.20.545792

**Authors:** Diandra Brkić, Sara Sommariva, Anna-Lisa Schuler, Annalisa Pascarella, Paolo Belardinelli, Silvia L. Isabella, Giovanni Di Pino, Sara Zago, Giulio Ferrazzi, Javier Rasero, Giorgio Arcara, Daniele Marinazzo, Giovanni Pellegrino

**Affiliations:** IRCCS San Camillo, Venice, Italy; Dipartimento di Matematica, Università di Genova, Genova, Italy; Max Planck Institute for Human Cognitive and Brain Sciences, Leipzig, Germany; Istituto per le Applicazioni del Calcolo “M. Picone”, National Research Council, Rome, Italy; CIMeC, University of Trento, Trento, Italy; Research Unit of Neurophysiology and Neuroengineering of Human-Technology Interaction (NeXTlab), Università Campus Bio-Medico di Roma, Rome, Italy; Philips Healthcare, Milan, Italy; CoAx Lab, Carnegie Mellon University, Pittsburgh, USA; Faculty of Psychology and Educational Sciences, Department of Data Analysis, University of Ghent, Ghent, Belgium; Department of Clinical Neurological Sciences, Western University, London, Ontario, Canada

## Abstract

Magnetoencephalography and electroencephalography (M/EEG) seed-based connectivity analysis requires the extraction of measures from regions of interest (ROI). M/EEG ROI-derived source activity can be treated in different ways. It is possible, for instance, to average each ROI’s time series prior to calculating connectivity measures. Alternatively, one can compute connectivity maps for each element of the ROI prior to dimensionality reduction to obtain a single map. The impact of these different strategies on connectivity results is still unclear.

Here, we address this question within a large MEG resting state cohort (N=113) and within simulated data. We consider 68 ROIs (Desikan-Kiliany atlas), two measures of connectivity (phase locking value-PLV, and its imaginary counterpart- ciPLV), three frequency bands (theta 4-8 Hz, alpha 9-12 Hz, beta 15-30 Hz). We compare four extraction methods: (i) mean, or (ii) PCA of the activity within the seed or ROI *before* computing connectivity, map of the (iii) average, or (iv) maximum connectivity *after* computing connectivity for each element of the seed. Hierarchical clustering in then applied to compare connectivity outputs across multiple strategies, followed by direct contrasts across extraction methods. Finally, the results are validated by using a set of realistic simulations.

We show that ROI-based connectivity maps vary remarkably across strategies in terms of connectivity magnitude and spatial distribution. Dimensionality reduction procedures conducted *after* computing connectivity are more similar to each-other, while PCA before approach is the most dissimilar to other approaches. Although differences across methods are consistent across frequency bands, they are influenced by the connectivity metric and ROI size. Greater differences were observed for ciPLV than PLV, and in larger ROIs. Realistic simulations confirmed that *after* aggregation procedures are generally more accurate but have lower specificity (higher rate of false positive connections). Though computationally demanding, *after* dimensionality reduction strategies should be preferred when higher sensitivity is desired. Given the remarkable differences across aggregation procedures, caution is warranted in comparing results across studies applying different methods.

## Introduction

Technological advances in neuroimaging over the past three decades have allowed the study of brain connectivity, which has helped to understand the neural basis of healthy cognition and clinical disorders (Schnitzler & Gross, 2005). Today it is well-established that task-based or resting state functional connectivity patterns are markers of efficiency in cognitive processes, while disrupted connectivity patterns may suggest impaired functional brain circuits (Aydin et al., 2020; Baldassarre et al., 2016; Carter et al., 2009; Englot et al., 2015; Hawkins et al., 2020; Pellegrino et al., 2012, 2019; Pellegrino, Mecarelli, et al., 2018; Schuler et al., 2018, 2019; Schuler & Pellegrino, 2021). Among many neuroimaging modalities used to estimate brain connectivity, magnetoencephalography and electroencephalography (M/EEG) are particularly effective because they provide sub-millisecond temporal-resolution, direct monitoring of neural activity oscillating across multiple frequencies (Baillet, 2017), acquisitions in a noise-free environment, either seated, laying down, or walking (Colenbier et al., 2022; Pellegrino et al., 2022; Schuler & Pellegrino, 2021).

While brain functions are spatially distributed, in order to measure connectivity the cerebral cortex is typically divided into parcels, regions of interest (ROI), or seeds, often standardized using atlases (Desikan et al., 2006; Eickhoff et al., 2018; Schaefer et al., 2018). Here, connectivity can be computed considering these regions in a seed-based connectivity fashion. Within this approach measures of functional relationship are typically estimated between one seed and other seeds, or between one seed and the rest of the brain, (Betti et al., 2013; Brookes et al., 2011; Siems et al., 2016). When computing seed-based connectivity, the seed can be a single voxel (for volumes) or a vertex (for analyses restricted to cortical surfaces), (Brookes et al., 2011; Hipp et al., 2012; Siems et al., 2016). More often seeds include larger regions consisting of multiple elements (voxels or vertices), containing rich and spatially varying information (Kriegeskorte & Bandettini, 2007; Meier et al., 2008). As each element has different connectivity, aggregation procedures are necessary in order to estimate the connectivity of a given region. One of the simplest dimensionality reduction strategies is to consider the average time-course of all elements of the ROI, which is standard procedure in fMRI seed-based connectivity (Basti et al., 2020; Brookes et al., 2011). This is reasonable given the spatial properties of the BOLD signal: large brain regions show some degree of homogeneity in signal and connectivity, allowing thus to extract brain parcellations (Fan et al., 2016; Schaefer et al., 2018; Thirion et al., 2014; Thomas Yeo et al., 2011).

In M/EEG the process of extracting seed-based connectivity is more challenging (Basti et al., 2019, 2020; Capilla et al., 2022; Farahibozorg et al., 2018; Hillebrand et al., 2012; Tait et al., 2021). M/EEG brain signals are not measured directly as the sensors are placed on the scalp (EEG) or several centimeters above (MEG). M/EEG brain activity is then reconstructed via source imaging, which requires resolving forward and inverse problems (Baillet, 2017; He et al., 2018; Pellegrino et al., 2020; Pellegrino, Hedrich, et al., 2016, 2018). M/EEG connectivity is then typically measured in this computed source space, as this reduces the bias due to volume conduction and signal leakage, and provides an accurate inference of the topology of brain connectivity (Brunner et al., 2016; Haufe et al., 2013; Hincapié et al., 2017; Lai et al., 2018; Palva et al., 2018; Schaworonkow & Nikulin, 2022; Van de Steen et al., 2019). When the source space is restricted to the cortical surface and the inverse solution is a distributed technique, source space time-courses are vectors of magnitude and direction for each vertex, that take into account cortical folding (Dale & Sereno, 1993; Hedrich et al., 2017). In other words, time-courses of neighboring vertices have different directions due to the curvature of the cortical folds. In these cases, time-course averaging within the parcel leads to a certain degree of signal cancellation (Ahlfors, Han, Belliveau, et al., 2010; Ahlfors, Han, Lin, et al., 2010; Chowdhury et al., 2018; Hillebrand et al., 2012; Irimia et al., 2012). Alternative approaches exist for extracting a single connectivity pattern from an extended seed, but the effects of these multiple strategies on connectivity patterns remain to be explored (Colclough et al., 2015, 2016; Hillebrand et al., 2012). For instance, one approach considers the time-course of the first PCA component rather than the average, as a representative of the ROI’s time course (Basti et al., 2020; Bruña & Pereda, 2021; Colclough et al., 2015, 2016; Dimitriadis et al., 2018). Another approach involves computing connectivity using the time-courses of all seed’s elements (voxel or vertex) and then average, PCA, or multivariate methods are performed (Aydin et al., 2020; Bruña et al., 2023, 2023; Colclough et al., 2016; Hillebrand et al., 2012; Palva et al., 2010). A ‘multivariate’ strategy acknowledges that all elements of the source space and their time-courses carry some useful information, while the aggregation approach such as mean or PCA is the easiest computational choice. Beyond these, a number of more complex procedures (identification of the center of mass, time-series with maximal power, weighted average, etc.) have been proposed, (Basti et al., 2019, 2020; Bruña et al., 2023; Chalas et al., 2022; Garcés et al., 2016; Korhonen et al., 2014; Luckhoo et al., 2012).

In short, the purpose of this study is to systematically compare the most common dimensionality reduction strategies to estimate connectivity patterns. Our analyses were performed using: (i) a large resting state MEG dataset (N=113) where we applied four extraction methods and two connectivity measures, across three frequency bands, using one canonical cortical atlas (Desikan et al., 2006). Furthermore we used realistic simulations in order to compare real data connectivity results with ground truth (simulated data).

Therefore, this work addresses the following questions: (i) How does the choice of ROI extraction method affect the estimation of resting state functional connectivity? (ii) Are differences between extraction strategies consistent across different connectivity measures (ciPLV, PLV) and frequency bands (theta, alpha, beta)? (iii) Does the reliability of extraction methods vary depending on the size of the ROI? (iv) What are the recommendations for future studies?

## 1. Methods

### 2.1 Participants and MEG acquisition

The study was approved by the local Ethics Committee, the Research Ethics Board of the Province of Venice (Italy), and complied with the 1964 Declaration of Helsinki and its later amendments. Every participant provided written and informed consent. All participants were healthy (self-reported) adults, with normal or corrected-to-normal vision, had no history of neuropsychiatric disorders, or brain injuries. Demographic details of the sample of participants included in this analysis are illustrated in Table S1 of the supplementary material. Overall, 84 subjects were included (66 Female, age range 21-58, mean 29 years old). Of these, 23 had two or more resting state sessions, in separate instances, that were treated as independent acquisitions. We analyzed 113 five-minute resting state recordings, acquired using a whole head 275-channel CTF system (VSM MedTech Systems Inc., Coquitlam, BC, Canada). Participants sat with their eyes closed in a magnetically shielded room. Eye movements (EOG) and cardiac rhythm (ECG) were recorded with bipolar electrodes. The sampling rate was set to 1200 Hz. Prior to the MEG data acquisition, the position of scalp points and anatomical landmarks (nasion, left and right pre-auricular points) were digitized with a 3D Fastrack Digitizer (Polhemus, Colchester, Vermont, USA). The head position within the dewar was tracked using the Continuous Head Localization system.

### 2.2 MRI data acquisition and analysis

All participants obtained MR images of the head after the MEG session. T1-weighted anatomical images were acquired at 1.5 T with the Achieva Philips scanner (Philips Medical Systems Best, The Netherlands) using the following parameters: TR = 8.3 ms, echo time TE = 4.1 ms, flip angle = 8°, isotropic spatial resolution = 0.87 mm.

### 2.3 MEG preprocessing and source imaging

All MEG data was analyzed with the Brainstorm MATLAB-based toolbox (Tadel et al., 2011) *(Figure 1)*. First, MRIs were imported and processed with CAT12 (Gaser et al., 2022) for tissue segmentation, cortical reconstruction and cortical labelling. Source space was defined as the cortical mesh extracted with CAT12, and downsampled to 4032 vertices. The ROIs were the 68 cortical regions of the Desikan-Killiany (DK) surface-based atlas (Desikan et al., 2006). MRI-MEG co-registration was performed by fitting a surface between the T1 head shape and the digitized head and fiducial points (acquired prior to the MEG recording, as described in our previous studies (Pellegrino, Machado, et al., 2016). MEG data was preprocessed using: spatial gradient noise cancellation of third order; band-pass filtering [0.3 – 256 Hz] and notch filtering (50, 100, 150 Hz); Signal Space Projection (SSP) to remove cardiac and eye movement artefacts (Taulu & Simola, 2006; Tesche et al., 1995); downsampling to 128 Hz; data segmentation into 2.5-second epochs; inspection and rejection of epochs affected by residual artefacts or head movement. The forward model was computed using overlapping spheres (Pellegrino, Hedrich, et al., 2018). Noise covariance was estimated from a 3-minute empty room MEG recording acquired prior to the experiment. The inverse problem was solved with the weighted minimum norm estimate (wMNE) approach, with dipoles at the source space constrained to be perpendicular to the cortical surface mesh (Brancaccio et al., 2020; Cona et al., 2020; Hämäläinen & Ilmoniemi, 1994; Hincapié et al., 2017; Pellegrino, Maran, et al., 2018).

**Figure 1.**
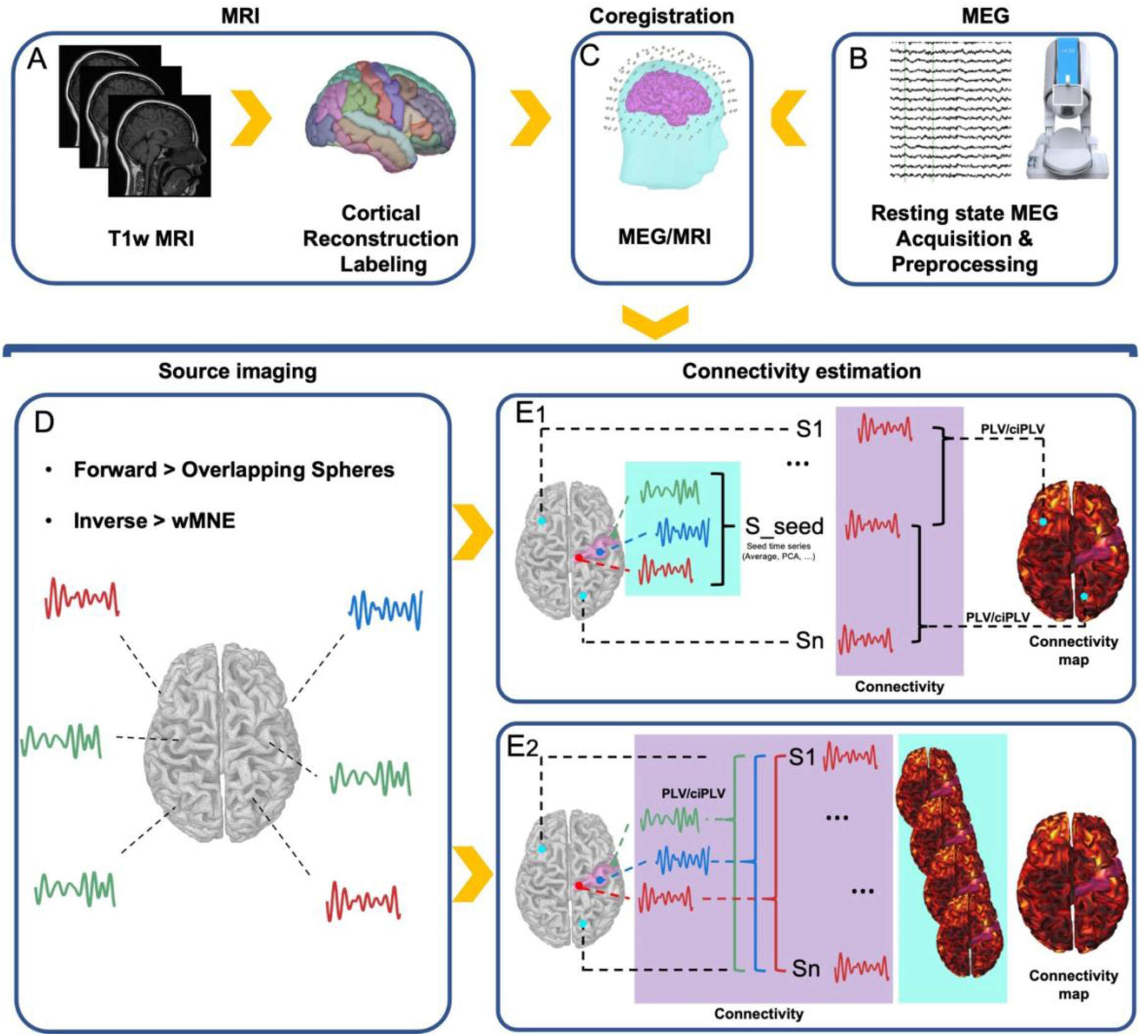
Data analysis pipeline. *<u>Panel A</u>* A volumetric T1w MRI was acquired for each participant and processed (segmentation, cortical reconstruction, cortical labeling) with CAT12.The source space corresponded to the cortical surface tasseled into 4000 vertices/nodes. *<u>Panel B</u>* MEG data was acquired with a 275 CTF system. Data underwent standard preprocessing. *<u>Panel C.</u>* MRI-MEG co-registration was performed with a surface fitting procedure taking into account fiducial points acquired with a Polhemus system *<u>Panel D.</u>* Source imaging was based on an overlapping sphere head model and wMNE inverse solution. This allowed reconstruction of time-series for each of the 4000 nodes of the source space. *<u>Panel E1.</u>* The dimensionality reduction procedure was applied to the time-courses of the vertices/nodes belonging to the ROI (cyan box). Two procedures were considered: (i) the average of the time courses and (ii) the PCA of the time-courses explaining the largest variance. Thereafter, the resulting time-course was considered as the seed and pairwise connectivity was estimated with each node/vertex of the source space (purple box). The result of this procedure was a single map representing the connectivity between the ROI and all nodes of the cortex. *<u>Panel E2.</u>* The dimensionality reduction procedure was applied to the connectivity maps computed for each vertex/node of the ROI and the source space (cyan box). The number of connectivity maps corresponds to the number of nodes/vertices of the ROI (purple box). The final output is a single map obtained as an average or maximum of all the maps previously computed.

### 2.4 Connectivity measures

Connectivity was computed with two measures of phase consistency: the Phase Locking Value (PLV; (Lachaux et al., 1999)) and its corrected imaginary counterpart, called corrected imaginary Phase Locking Value (ciPLV; (Bruña & Pereda, 2021)). PLV is a measure of phase consistency between two oscillating time series, where higher values represent higher connectivity. More specifically, PLV measures the instantaneous phase difference between two signals based on the assumption that the phase of connected signals are aligned and evolve together (Lachaux et al., 1999; Nolte et al., 2020; Varela et al., 2001). It has been shown how the application of the imaginary part of the complex definition of PLV, ciPLV, is less sensitive than PLV to volume conduction and signal leakage, since it does not consider zero-lag phase alignment (Bruña et al., 2018; Colclough et al., 2016; Palva et al., 2018; Schuler et al., 2022; Tabarelli et al., 2022). The mathematical explanation of differences between PLV and ciPLV are discussed in detail in (Nolte et al., 2020). We applied PLV and ciPLV in three canonical frequency bands of interest: theta (5-7Hz), alpha (8-12Hz), and beta (15-29Hz), (Bruña et al., 2018; Nolte et al., 2020). Connectivity was estimated for each frequency band, each of PLV and ciPLV, and between each ROI (68 ROIs of the Desikan-Kiliani atlas) and each element of the source space, resulting in a connectivity map. Each of these maps was represented by an array of 4032 elements, where the value of each element is the connectivity between the ROI and each element (vertex) of the source space. In order to perform group analyses, individual connectivity maps were projected onto a common template (MRI-ICBM152) (Mazziotta et al., 2001) and smoothed with a full width at half maximum of 5mm (Bernal-Rusiel et al., 2010; Brodoehl et al., 2020; Coalson et al., 2018; Hagler et al., 2006; Worsley et al., 2002).

### 2.5 Dimensionality reduction methods or ROI extraction function

We compared four methods of ROI extraction, illustrated in Figure 1:

a. *Mean before:* mean of the signal within ROI. This function averages all time-courses within the ROI before computing connectivity between the resulting average time-course and the time-course of each element of the source space. This approach is very simple and computationally efficient. As the time-course of adjacent elements of the source space may display a different sign due to the folding of the cortical surface, a flip-sign function is applied before averaging in order to reduce cancellation.
b. *PCA before:* PCA of the signal within the ROI. This method takes the time course of the first component of the PCA decomposition of all the signals within a ROI, before computing the connectivity between the time course of that component and the time course of all elements of the source space.
c. *Mean after:* mean of the connectivity maps computed from all elements of a ROI. Here, dimensionality reduction is applied after the computation of connectivity. A connectivity map is computed for each element of the ROI taking into account the time course of that element and the time-course of all elements of the source space. The final connectivity map corresponds to the average of all the maps belonging to the same ROI.
d. *Max after:* maximum estimate of the connectivity computed for all elements of the ROI. It is similar to the mean after, but in the Max after function only the maximum value is retained for each element of the resulting connectivity map/vector.

### 2.6 Clustering analysis

We applied hierarchical clustering analysis to examine the similarities between the aforementioned extraction methods, and two connectivity measures, PLV and ciPLV, across three frequency bands (theta, alpha, and beta). Specifically, for each frequency band we first computed the Pearson’s correlation coefficient between connectivity maps resulting from the combination of connectivity measure and scout function. This resulted in an 8 x 8 similarity matrix R for each dataset. Subsequently, a distance matrix was calculated as 1 - <R>, where <R> denotes the data-averaged similarity matrix. Finally, a hierarchical clustering was applied to this distance matrix using the MATLAB linkage function and selecting the average linkage criterion. Importantly, we performed this clustering analysis at two different brain granularity levels: entire cortical surface and for each ROI separately. For the former, connectivity maps were concatenated across all seed regions (68 ROI of the DK atlas). For the latter, we considered ROI-specific maps. Lastly, to explore whether the patterns observed were frequency-specific, we looked at the average distribution across three frequency bands of interest, theta (4-8Hz), alpha (8-12Hz), and beta (12-25Hz). This entire clustering analysis pipeline is openly available on <u>Github</u>, (jrasero, 2022/2023).

### 2.7 Pairwise comparison between scout functions

While the clustering analysis provided an estimate on similarity/dissimilarities across different strategies, and how connectivity maps cluster together, it did not quantify differences across options and their spatial distribution. Therefore, in order to address the extent to which the use of the four scout functions differed, we performed parametric testing. Specifically, we applied paired t-testing to compare across all possible combinations of scout functions, for each ROI, frequency bands and connectivity measures (PLV and ciPLV). This analysis was carried out as a post-hoc exploration of the magnitude and spatial distribution of differences across the dimension reduction approaches. We found that the differences across procedures were strong and widespread. Therefore, in order to make sure that these differences were specific and not caused by a simple offset (i.e. a constant difference), we repeated the same analysis after normalizing the connectivity maps. To this end, the maximal connectivity value of each map was set as 100% and all other values were expressed as percentages of the maximum. For these analyses we only report some examples, whereas all comparisons are available as GIfTI at www.hsancamillo.it.

### 2.8 Simulation of MEG data with known connectivity structure

Applying different ROI-extraction methods to realistic simulations allowed us to depict the properties of each approach in comparison with the ‘ground truth’. The MEG forward model was based on individual anatomical data of one of the participants. Consistently with the analysis of real data, the source space was confined to the cortical surface, and the lead field matrix was computed by applying the overlapping sphere model. We simulated neural activity by mimicking five interacting patches (for a schematic representation of the implemented pipeline, refer to Figure 2). For each ROI we randomly drew a point, *v1*, within the ROI and four other vertices, *v2, v3, v4, v5,* outside the selected ROI, with the only constraint being that the distance between each pair of points was at least 5 cm. Around each point *vi*, *i=1,…,5*, we defined a patch *Pi* by considering *vi*’s neighbouring source-space elements that were less than 2.5 cm apart. Additionally, as for *P1,* we restricted it to entirely belong to the ROI containing *v1* and we set three possible sizes of the patch, namely (i) all points with a distance from *v1* lower than 2.5 cm, (ii) 50% of those points by selecting the closest to *v1*, and (iii) only *v1* (Figure 2, Panel A). For each group of patch configurations, we then simulated three time-courses, s1(t), s2(t), s3(t), t=1, …., T, where *T=10000* mimicking about 78 s of neural activity sampled at 128 Hz. To this end, and similar to previous work (Haufe & Ewald, 2019; Sommariva et al., 2019; Vallarino et al., 2021), only alpha band was considered for simulation using three signals following a multivariate autoregressive (MVAR) model of order 5, so that *s1(t)* leads the activity of *s2(t)*, while *s3(t)* is uncorrelated. We only retained stable MVAR models such that (i) for each of the resulting signals the average power spectrum in the alpha band represented at least 40% of the overall average power spectrum, and (ii) the average coherence in the alpha band between *s1(t)* and *s2(t)* was greater than 0.5. Following this, *s1(t)* was assigned to *v1*, *s2(t)* to *v2* and *v4*, and *s3(t)* to *v3* and *v5*. For each patch *Pi* the activity of the remaining points was defined by randomly perturbing the Fourier transform of the time series of the corresponding centre *vi* so as to reach a certain amount of intra-patch coherence (Hincapié et al., 2017). Additionally, a Gaussian window was used to modulate the resulting time-series so that source intensity decreased for increasing distance from *vi* (Figure 2, Panel B). Finally, for each set of patch activities, we computed the magnetic field at sensors and added simulated additive noise according to the random dipole brain noise model PoMAM (Calvetti et al., 2019; de Munck et al., 1992) (Figure 2, Panel C). Hence a total of N=68×3=204 MEG data points were simulated, 68 being the number of ROIs within the Desikan-Kiliany atlas and 3 being the considered sizes for *P1*.

**Figure 2.**
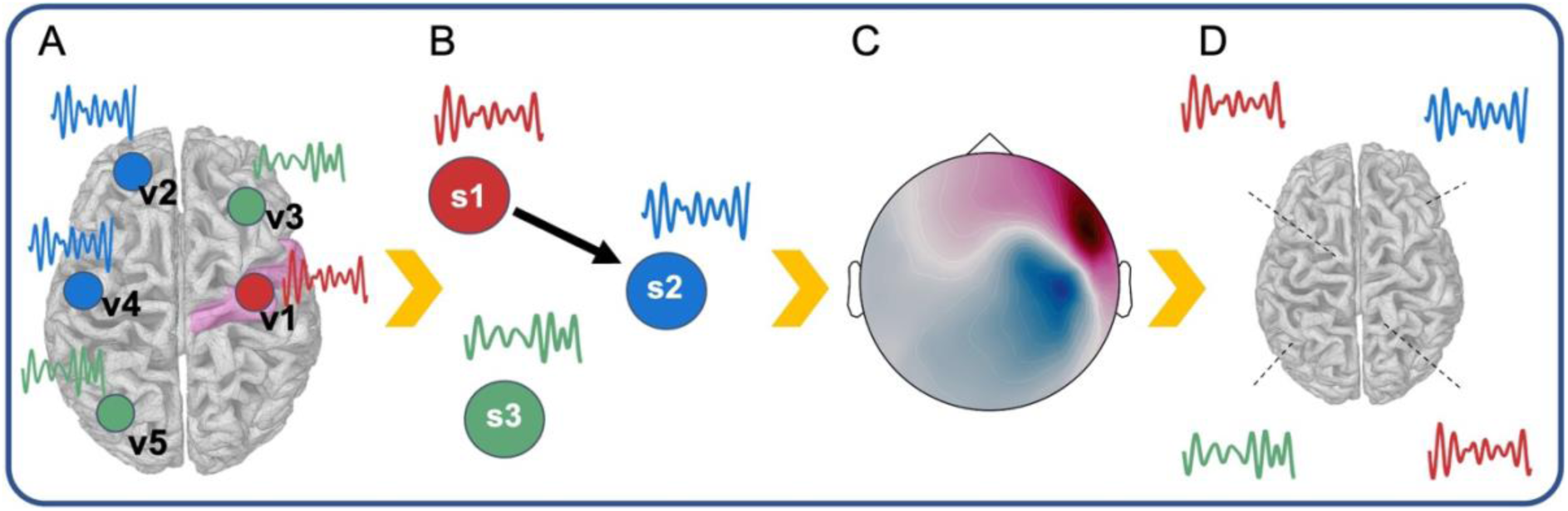
Simulation Pipeline. *<u>Panel A.</u>* For each ROI we randomly drew a point, v1, within the ROI and four other vertices, v2, v3, v4, v5, outside the selected ROI, with the only constraint that the distance between each pair of points was at least 5 cm. Around each point vi, i=1,…,5, we defined a patch Pi by considering vi’s neighbouring source-space elements that were less than 2.5 cm apart. We set three possible sizes of the patch, namely (i) all points with a distance from v1 lower than 2.5cm, (ii) 50% of those points by selecting the closest to v1, and (iii) only v1. *<u>Panel B</u>*. For each group of patch configurations, we simulated three time-courses, s1(t), s2(t), s3(t) so that s1(t) leads the activity of s2(t), while s3(t) is uncorrelated. s1(t) was assigned to v1, s2(t) to v2 and v4, and s3(t) to v3 and v5. For each patch Pi the activity of the remaining points was defined by randomly perturbing the Fourier transform of the time series of the corresponding centre vi so as to reach a certain amount of intra-patch coherence. *<u>Panel C.</u>* for each set of patch activities, we computed the magnetic field at sensors and added simulated additive noise according to the random dipole brain noise model PoMAM *<u>Panel D.</u>* A total of N=204 MEG data points were simulated. Source space signals were reconstructed using a similar procedure as was used for real data.

### 2.9 Connectivity estimate and evaluation criteria

As with the experimental MEG data, neural activity was estimated from the simulated MEG data by using the wMNE inverse solution, while connectivity was quantified from the source space estimated time-courses through PLV and ciPLV. Specifically, for each of the 204 simulated signals and for the two connectivity measures, we estimated cortical connectivity maps by considering as seed the ROI containing *v1*. The four ROI extraction methods, i.e. Mean before, PCA before, mean after, and Max after, were used to quantify the connectivity between this ROI and all 4002 other elements of the source space (Figure 2, Panel D).

To evaluate the results, we exploited the fact that, when P1 only includes *v1*, a ground truth can be defined by computing the values of PLV and ciPLV between *s1* and the activity in all the other points of the source space. Hence for these simulated MEG data, connectivity metrics, and scout function, we computed the Pearson correlation coefficient between the estimated cortical connectivity maps and the corresponding ground truth. On the other hand, when *P1* includes more than one element, defining a ground truth is not straightforward. For this reason, we also evaluated the accuracy of the estimated connectivity maps by computing true and false positive rates (TPR and FPR, respectively). To this end, we fixed one hundred thresholds uniformly distributed between 0 and 1 (alpha). Then for each of the simulated MEG data points, for each of the four scout functions and for both connectivity measures, we applied a normalization procedure by rescaling each map with its maximum value. Thereafter, for each map, for every alpha value of the threshold, we defined

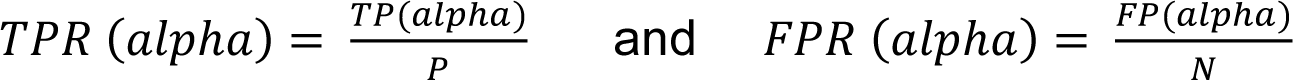

where:

- *P* and *N* are the number of positive and negative, respectively. *P* counts the source-space points truly connected with *v1*, i.e. the nodes within *P2* and *P4* in our simulations, while *N* is the number of the remaining source-space points without considering the nodes of the patch *P1* centered in *v1*.
- *TP(alpha)* is the number of true positives, i.e. points of the source space truly connected to *v1* where the normalized reconstructed connectivity value (either PLV or ciPLV) exceeded the threshold alpha.
- *FP(alpha)* is the number of false positives, which are points not truly correlated with *v1* but where the normalized reconstructed connectivity still exceeded alpha.

Finally, results in terms of True/False positive rate were summarized by computing the corresponding *Receiver Operating Characteristic (ROC)* curve and the associated *Area Under the Curve (AUC)*.

## 2. Results

### 3.1 Hierarchical Clustering

The results of the hierarchical clustering are summarized in *Figure 3*. This analysis showed similar patterns of similarity-dissimilarity across frequency bands *(Figure 3, rows)* and the two connectivity measures of interest *(PLV and ciPLV)*.

**Figure 3.**
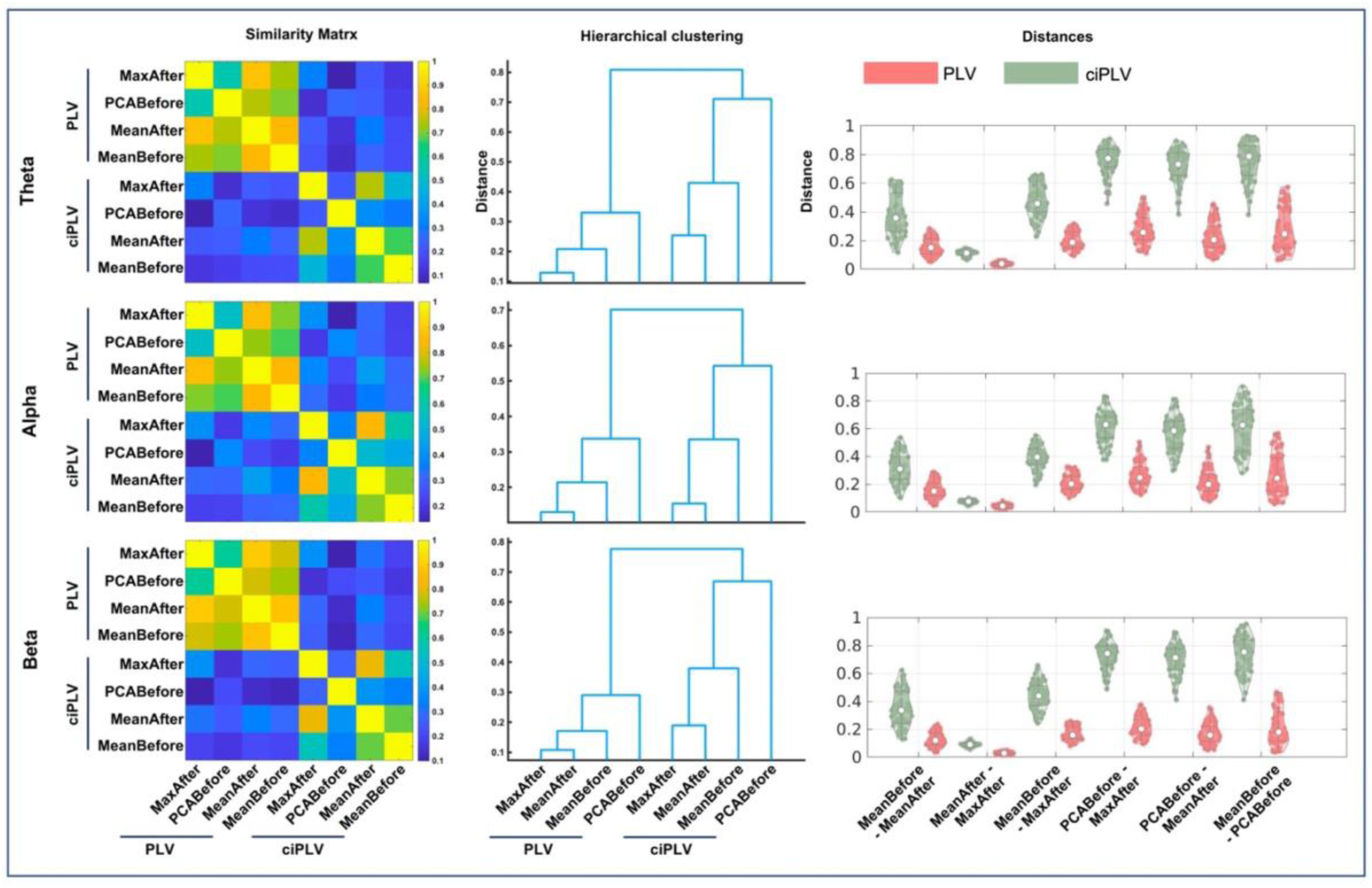
Hierarchical Clustering. The figure shows the clustering results by *<u>frequency band</u>* (*<u>rows</u>*). *<u>Left column:</u>* similarity matrices. The color scale indicates the similarity of connectivity maps across connectivity measures (*PLV and ciPLV*) and extraction methods (*Max after, PCA before, Mean after, Before*). *<u>Middle column:</u>* Dendrograms with distances across aggregation procedures. *<u>Right column:</u>* Distances across extraction procedures by connectivity measure (*PLV* and *ciPLV*). Map similarity was overall higher and distances overall lower for *PLV* than *ciPLV (Left and right columns). PLV* and *ciPLV* maps cluster together within connectivity measures for all frequency bands *(middle column)*. The relative distances across extraction methods also clustered consistently together within connectivity measure (*PLV* and *ciPLV*, *middle column*). The most similar extraction procedures were *Max after* and *Mean after*, which were closely clustered, followed by Before, which clustered with both *Max after* and *Mean after*. *PCA before* was most distant from all other approaches, as seen in both *middle and right columns*. The right column shows that *Mean after* and *Max after* had the lowest distance, followed by *Before*-*Mean after*.

Similarity map was overall higher and distances overall lower for PLV than ciPLV *(Figure 3, Left and right columns). PLV* and *ciPLV* maps clustered together within connectivity measures (*PLV and ciPLV*), for all frequency bands *(Figure 3, middle column)*. The relative distances across extraction methods also clustered consistently together within connectivity measure (*Figure 3, middle column*). We observed that *Max after* and *Mean after* were the two most similar extraction procedures, and as such clustered together under all circumstances (for all frequency bands and connectivity measures, *Figure 3, middle column*). *Max after* and *Mean after* were relatively close to *Before*, and clustered together, whereas *PCA before* had higher distance from all other approaches *(Figure3, middle and right columns)*.

The right column demonstrates that *Mean after* and *Max after* had the lowest distance, followed by *Before-Mean after*.

The topographical distribution of the distances across extraction methods is highlighted in *Figure 4*, left panel. The lowest distances were found in the deep regions especially of the midline, for all frequency bands. The higher distances were found especially in the convexity, over the fronto-central regions, and parietal regions. *Figure 4*.

**Figure 4.**
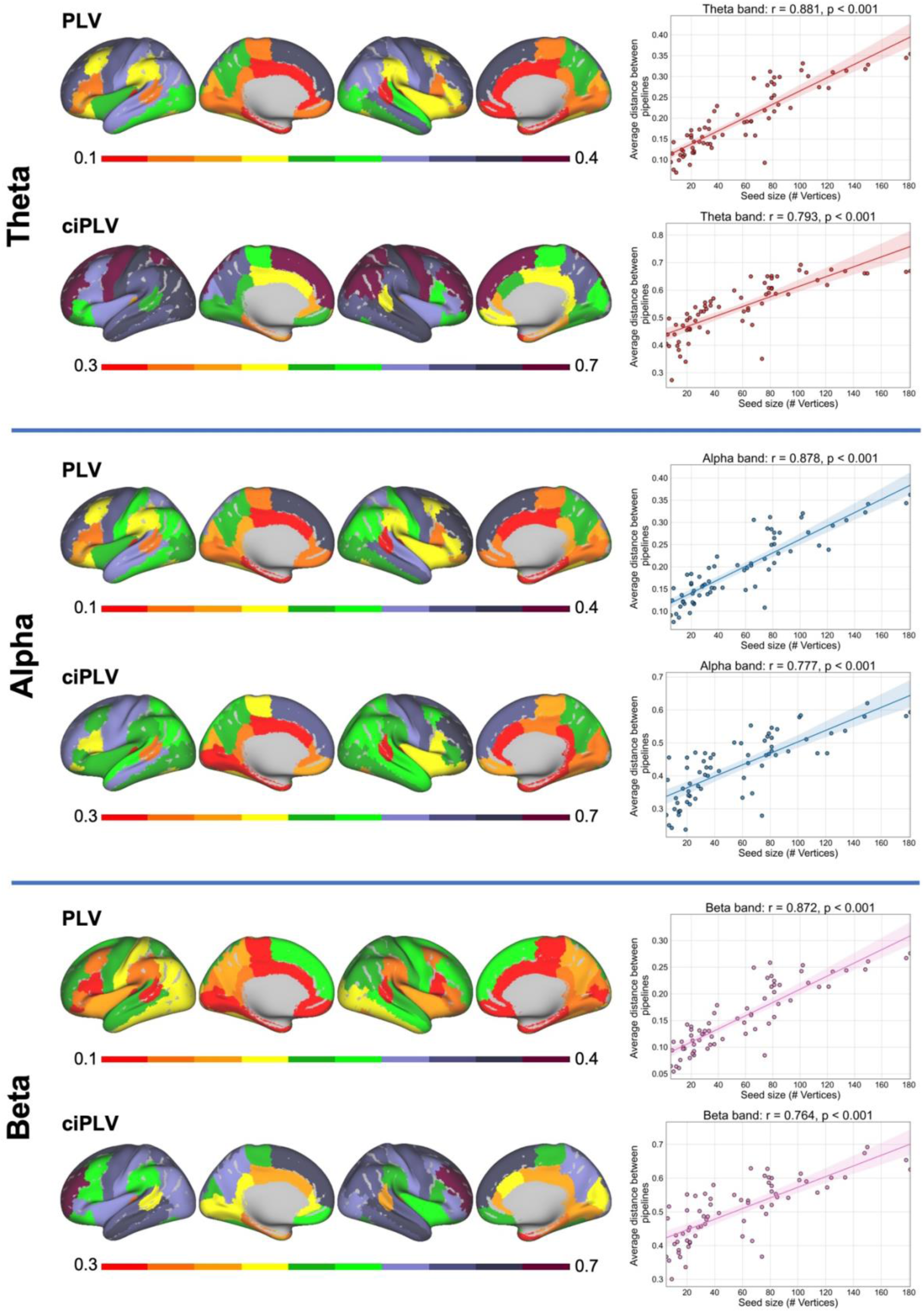
Topographical distribution of average distances by frequency band (rows) and relationship between the size of the parcel and average distances. *<u>Left Panel</u>* The lowest distances were found in deep regions, especially of the midline, across connectivity measures and frequency bands. The higher distances were found especially in the convexity, over the fronto-central regions, and parietal regions. *<u>Right</u> <u>panel</u>* There was a significant positive relationship between the size of the region and the average distance across extraction methods.

There was a significant and positive relationship between the size of the region and the average distance across extraction methods (*Figure 4, Right Panel*) (*Pearson’s correlation coefficient; r>0.7 and p<0.001 consistently across frequency bands and connectivity measures)*.

### 3.2 Paired comparisons

The cluster analysis provides a pattern of similarities but does not provide an immediate estimation of the magnitude and spatial distribution of differences across strategies. Therefore, we followed up on previous analysis by estimating how connectivity changed by paired comparisons between the four strategies under investigation. Specifically, for each connectivity measure, and within each frequency band, we compared: i) *Mean before* vs *PCA before*; ii) *Mean before* vs *Mean after*; iii) *Mean before* vs *Max after*; iv) *PCA Before* vs *Mean after*; v) *PCA Before* vs *Max after*; vi) and *Mean after* vs *Max after*. For each of these comparisons, we computed t-maps as a representation of normalized differences between the two strategies (*Figure 5*). ciPLV connectivity maps computed showed differences as high as 40 t-values, with magnitude and direction (positive/negative) were consistent across frequency bands (alpha, beta, theta). Similar patterns were observed for PLV (*supplementary material*).

**Figure 5.**
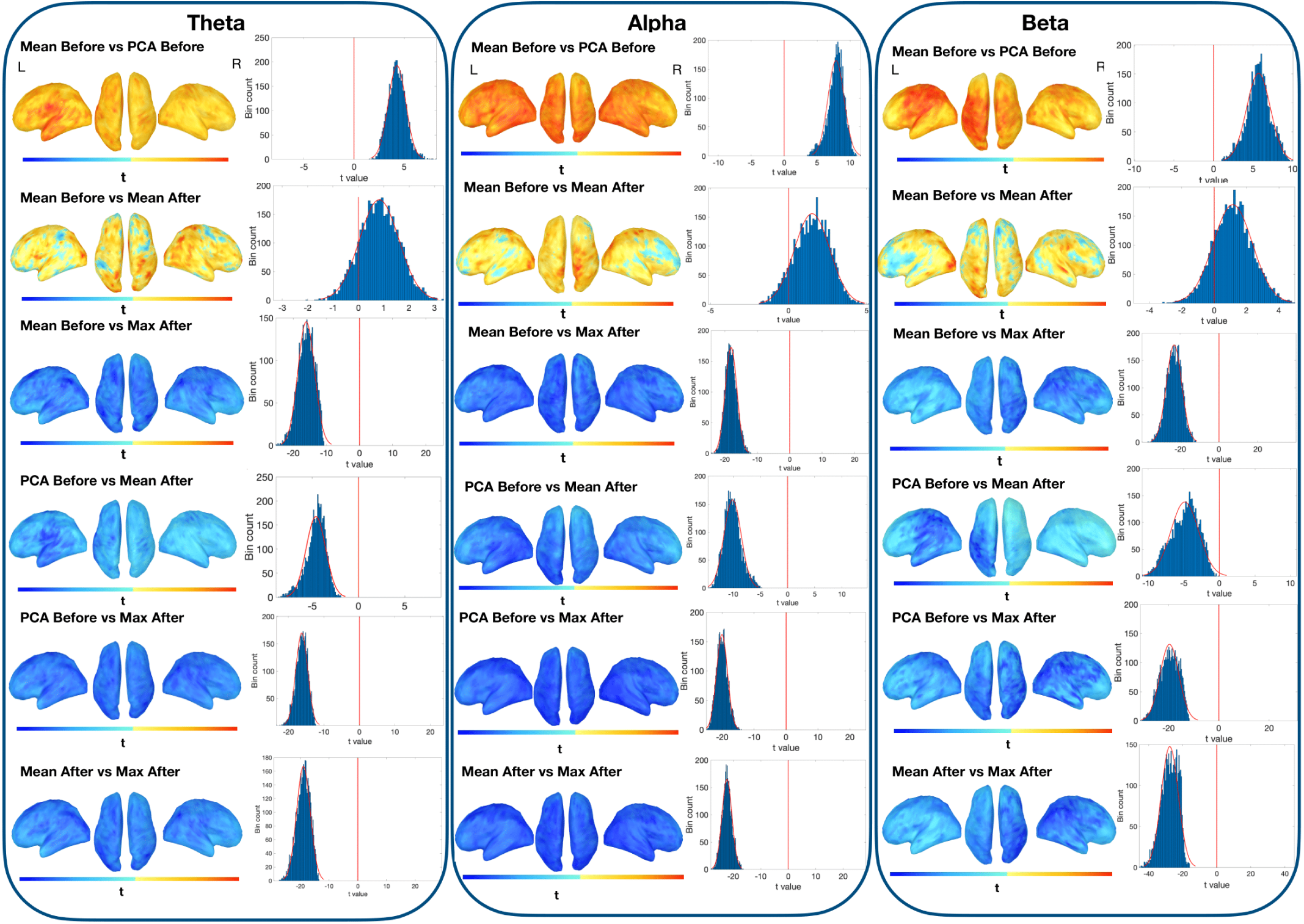
ciPLV t-maps of pairwise comparisons between connectivity maps of the left superior temporal gyrus, across aggregation procedures, for theta, alpha and beta bands. Color scale is adjusted for each comparison and ranges between [-max to +max] t-value. Histograms show t-value distribution across the entire cortical surface.

### 3.3 Simulations results

#### 3.3.1 Correlation between true (simulated) and estimated maps

The results of the correlation between connectivity estimated and true (simulated) when the patch P1 contains only its center v1 are summarized in *Figure 6*. For both PLV and ciPLV and for all aggregation procedures the correlation coefficients are rather low *(< 0.15, Figure 6, upper row)*. This result is expected, as the simulated map does not contain any activity for a large portion of the cortical surface, whereas the estimated map may contain some activity over the entire surface as a result of signal leakage associated with the inverse solution. Similarly, it is expected that PLV provides lower values for this correlation, since it is more sensitive to source leakage than ciPLV. Finally, negative values of the correlation coefficient correspond to data where the value of connectivity estimated in the patches that were uncorrelated with v1, namely P3 and P5, was on average higher than the value of connectivity estimated in the correlated patches, namely P2 and P4. In this scenario, the aggregation approaches applied after computing connectivity generally perform better. When applying PLV, the correlation between estimated and simulated connectivity is significantly higher (better) for *Mean after* as compared to *Mean before* and *PCA before* (*Wilcoxon signed-rank, p<0.05 consistently*), whereas when applying ciPLV Max after performs slightly but significantly better than all other approaches (*Max after vs PCA before W=1761, p=0.003; Max after vs Mean before W=1897, p=0.0001; Max after vs Mean after W=1690, p=0.02*) and Mean after performs slightly better than Mean before (*W =1793, p=0.002*).

**Figure 6.**
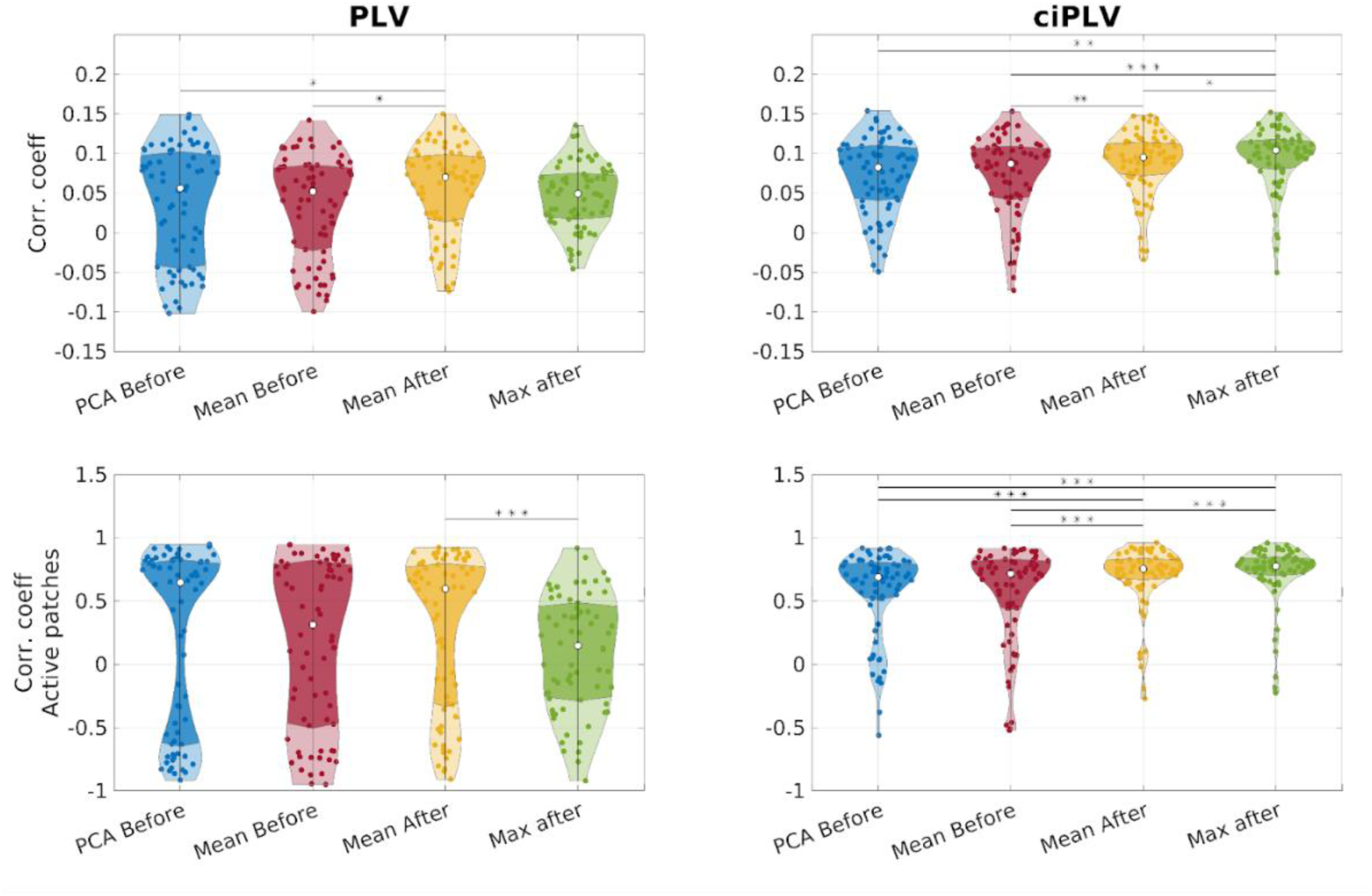
Pearson correlation coefficient between true (simulated) and estimated connectivity maps across 68 simulated neural activity sources such that P1 contains only the center v1. *<u>Upper row, left panel.</u>* PLV correlation coefficients of maps defined in the entire source-space. *Mean after* performed better than *Mean before* and *PCA before*. *<u>Upper row, right panel</u>* ciPLV map correlation coefficients defined for the entire source-space. *Max after* performed significantly better than all other aggregation approaches. *Mean after* performed significantly better than *Mean before*. *<u>Lower row,</u> <u>left Panel</u>* PLV correlation coefficients of maps containing only P2-P5. The distribution of correlation coefficients was remarkably variable across aggregation procedures. *Mean after* performed significantly better than *Max after. <u>Lower row, Right Panel</u>* ciPLV correlation coefficients of maps containing only P2-P5. Both *Mean after* and *Max after* performed significantly better than *Mean before* and *PCA before.* Note, in the violin plots, white dots depict median values. Darker colors correspond to the interquartile range. ∗ p < 0.05, ∗∗ p < 0.005, ∗∗∗ p < 0.0005.

We repeated the correlation analysis by restricting true and estimated connectivity maps to the points pertaining to the four active patches, namely P2-P5 (*Figure 6, bottom row*). In this scenario, correlation coefficients are remarkably higher, especially for ciPLV. The aggregation approaches applied after computing connectivity remained slightly better, especially for ciPLV. The correlation coefficient of both ciPLV Mean after and ciPLV Max after were significantly higher than ciPLV Mean before and ciPLV PCA before (*Mean after vs Mean before W=1944, p=3*10^-5; Mean after vs PCA before W=2023, p=2.5*10^-6; Max after vs Mean before W=2043, p=1.3*10^-6; Max after vs PCA before W=2018, p=2.9*10^-6*). Note that no statistically significant difference was observed comparing ciPLV Max after and Mean after. Interestingly, in this scenario the correlation coefficient was less stable for PLV, with high levels of variance depending on the aggregation procedure. Here the only significant difference was in the comparison between Mean after which was higher than Max after *(W=1835 p=6*10^-4)*.

#### 3.3.2 Accuracy of estimated maps (true and false positive rates)

The analysis of False/True Positive rates provides more insight in interpreting results from the clustering analysis performed on the experimental resting-state data. *FPR(alpha)* and *TPR(alpha)* are reported in *Figure 7*. This analysis is focused on the case of P1 only containing the center v1, which is a scenario analogous to that applied in section 3.3.1.

**Figure 7.**
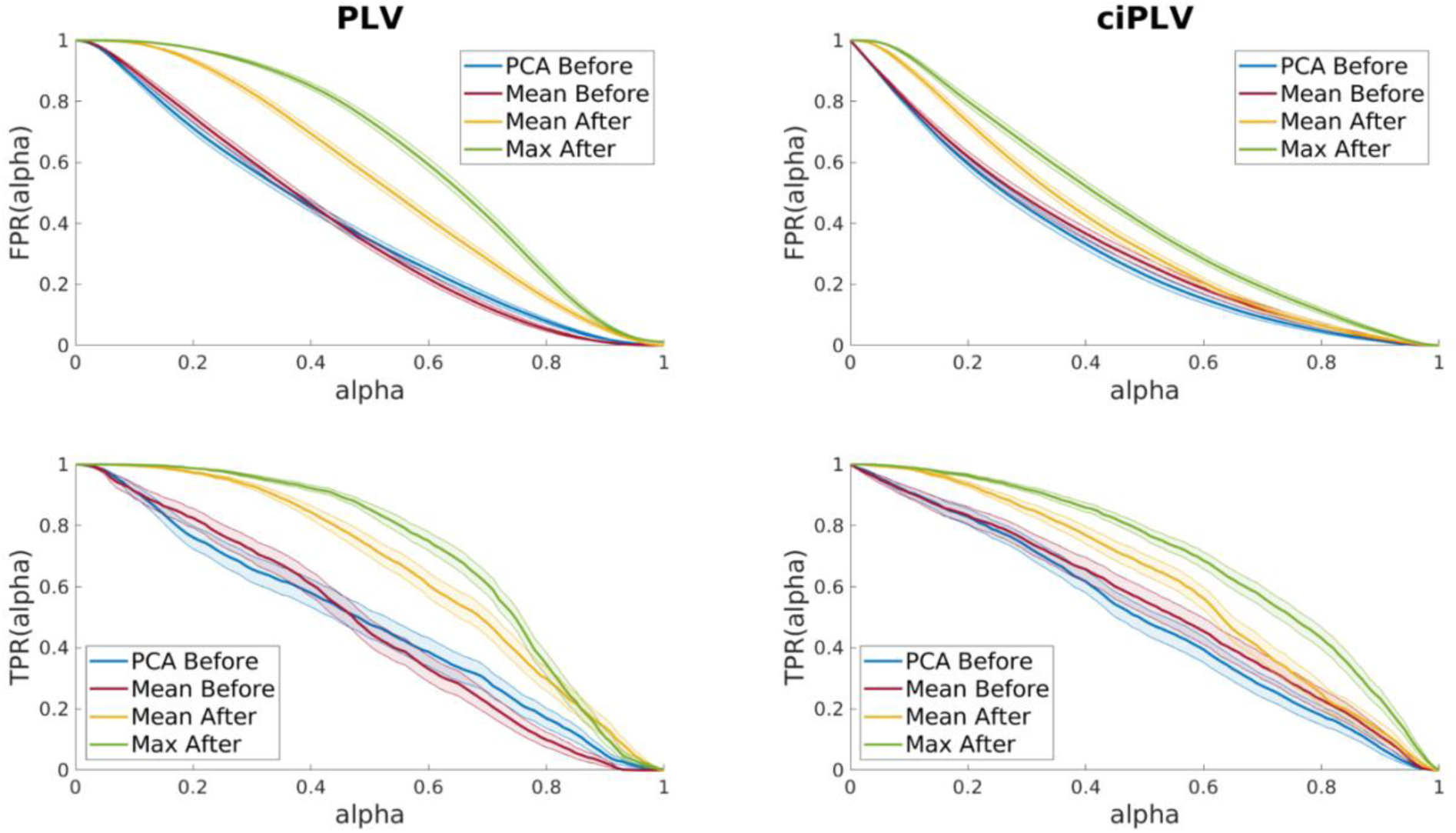
False Positive Rates (*upper row*) and True Positive Rates (*lower row*) models of the two connectivity measures (PLV -*left*- and ciPLV -*right*-) extracted via the four different aggregation procedures. Plots show mean and standard error of the mean across 68 simulated neural activity so that P1 only contains the centre v1. Results concerning different sizes of P1 can be found in the Supplementary Materials.

For both PLV and ciPLV the two *before* procedures *(Mean before and PCA before)* show higher specificity, while the two *after* procedures *(Mean after and Max after)* show higher sensitivity. Indeed, *PCA before and Mean before* show a lower false positive with the tradeoff of identifying fewer true connections. In other words, these aggregation procedures seem to be more conservative, in the sense that the reconstructed connections most likely identify truly connected sources, at the expense of weak connections which may be lost when these aggregation procedures are used. Vice versa, *Mean after* and *Max after* show a higher value of both true and false positive rates. This seems to suggest that weak connections may be retrieved at the expense of retaining spurious connectivity. Note also that for both ciPLV and PLV the FPR and TPR curves are very similar and overlap for most of the alpha range.

Similar results are obtained when varying the size of PI *(supplementary material)*. Although the differences across aggregation procedures can vary, Max after and Mean after typically show a higher rate of FPR and TPR.

The comparison of the AUC for all conditions under investigation *(Figure 8)* shows a significantly higher value for Max after and Mean after for ciPLV when only the centre of P1 is considered (*Mean after vs Mean before W=1792, p=0.002; Mean after vs PCA before W=1830, p=7*10^-4; Max after vs Mean before W=1914, p=7.15*10^5; Max after vs PCA before W=1907, p=8.75*10^-5*). The distribution of the AUC values is similar for the four ROI-extraction methods when all points of the patch P1 are considered. With respect to the PLV metric, the distribution of AUC values is significantly higher for Mean after regardless of the dimension of the P1 patch (*Mean after vs PCA before W=1656, p=0.037 when P1 containing only v1; Mean after vs Max after W=2068, p=5.4*10^-7 when P1 contains half points; Mean after vs Max after W=1957, p=2*10^-5 when P1 contains all voxels*).

**Figure 8.**
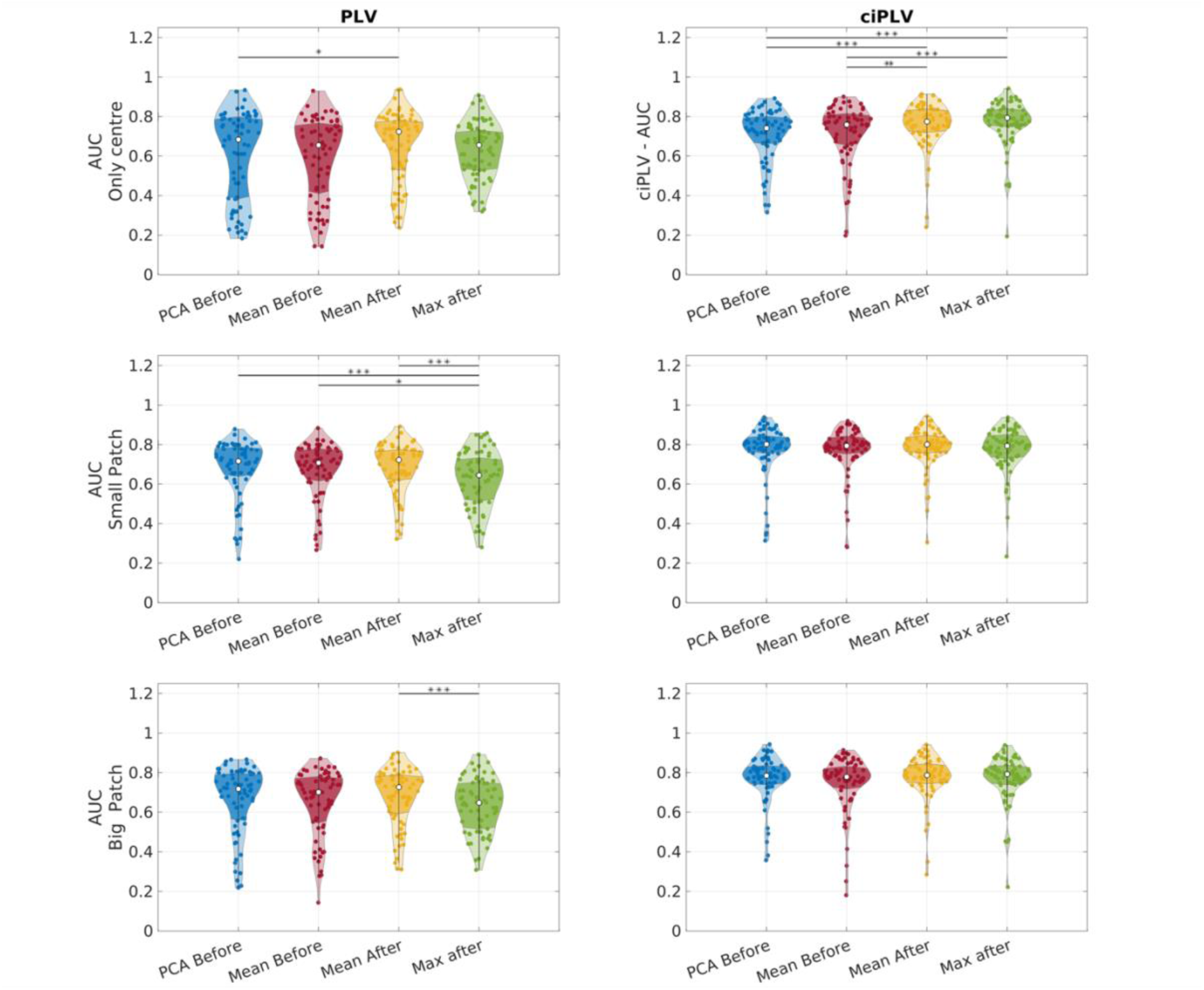
Distribution of the AUC values across the simulated MEG data when the patch P1 contains all the points that are less than 2.5cm apart from v1 (third row), half of such points (*second row*), only its centre v1 (*first row*). Left column shows results concerning PLV while right column refers to ciPLV.

## 3. Discussion

The findings of this study indicate that the choice of dimensionality reduction method has a significant impact on the resulting connectivity outcomes. These results were observed across both resting state data and realistic simulations. Interestingly, the choice of extraction method resulted in considerable differences in both the magnitude and distribution of connectivity. More specifically through the use of cluster analysis, we observed that the aggregation procedures applied after computing connectivity yielded more consistent results. This was observed for both in-vivo and simulated data.

The *after* approaches are computationally demanding, but allow for the integration of information from all elements of a given parcel, thus providing a more comprehensive connectivity estimation (Aydin et al., 2020; Bruña et al., 2023, 2023; Colclough et al., 2016; Hillebrand et al., 2012; Palva et al., 2010).

The results of the clustering analysis also revealed that the *Mean before* method produced connectivity maps that were similar to those generated by the *after* strategies. On the other hand, *PCA before* method yielded results that were markedly dissimilar from all other approaches. While cluster analysis alone may suggest that the *Max after*, *Mean after*, *Mean before* and *Mean after* methods produce similar results, it is important to note that the cluster approach is based on correlations across maps and does not provide enough information about the actual topographical distances. Therefore to gain a better understanding, we conducted pairwise analysis of connectivity maps generated by different extraction strategies. Our results revealed that the most similar maps differed by a magnitude of 40+ t-values, indicating that even minor variations in method choice lead to significant differences in connectivity outcomes (*Figure 5*). We also ruled out the possibility that these differences were due to an offset of connectivity values by directly contrasting normalized maps, where each connectivity value was rescaled against the maximum. Here we observed important and widespread differences in connectivity outcomes. Hence, caution is warranted when comparing connectivity patterns between studies that employ different extraction strategies, as even small variations in method can lead to significant discrepancies.

Our analysis also shed light on some of the factors that may influence the effect of extraction method on connectivity outcomes. Specifically, we observed that similarity across extraction methods was higher for *PLV* as compared to *ciPLV* (*Figure 3*). Both connectivity measures utilized in this study are based on phase synchrony across time series. However, *ciPLV* is insensitive to Lag=0 connectivity and as such, is less susceptible to volume conduction and spatial leakage (Bruña et al., 2018; Bruña & Pereda, 2021; Colclough et al., 2016; Lachaux et al., 1999; Nolte et al., 2020; Palva et al., 2018; Schuler et al., 2022; Tabarelli et al., 2022; Varela et al., 2001). Therefore, the resulting connectivity maps are typically less smooth, possibly explaining why similarities across extraction methods is lower than for *PLV (Figure 3)*. On the other hand, our results also highlight that other connectivity measures may exhibit varying levels of robustness to the extraction methods. We observed no relevant differences across frequency bands, although the spatial distribution of the distances shows a trend towards greater differences for lower frequencies (*Figure 3 and 4*). Low frequency activity recorded from the scalp typically originates from large populations of neurons or from large distribution of high frequency activity (Pellegrino et al., 2017; Tao et al., 2007). Therefore, it is possible that differences in aggregation methods are more pronounced when the underlying generators span widespread brain areas. This idea is supported by the significant positive linear relationship we observed between the size of the parcel (expressed in number of vertices) and the average difference across extraction methods and frequency bands (*Figure 3*).

*Figure 4* provides some insight into the topographical distribution of the differences. Smaller differences are more often observed in deep regions especially of the midline. MEG spatial sampling is not homogeneous across the cortical surface, and the reconstruction of source signal in deep, basal, and midline regions is not always accurate in MEG (Ahlfors, Han, Belliveau, et al., 2010; Gavaret et al., 2014; Huiskamp et al., 2010). It is therefore possible that the estimation of cortical connectivity was less refined in those areas, and the differences across methods were smaller.

Alternatively, the distribution of rhythms across the cortical surface with more alpha activity in the posterior quadrant, more theta activity in the temporal lobes, and faster activity in the range of beta band in the central regions may also play a role in determining this spatial distribution. Overall, these findings suggest that the choice of extraction method may have a greater impact on connectivity measures in lower frequency bands and in regions that are more challenging to measure using MEG. While MEG resting-state data allowed us to identify relative differences across extraction methods, it was challenging to determine which method was more accurate. This prompted us to conduct realistic simulations to gain a more comprehensive understanding of the performance of different extraction methods (Grova et al., 2016; Hincapié et al., 2017; Machado et al., 2018; Sommariva et al., 2019).

Our simulations confirmed that aggregation procedures performed after the computation of connectivity may provide more accurate results. The correlation coefficient between simulated and estimated maps of connectivity was significantly higher for *Mean after* for *PLV*, and for *Mean after* and *Max after* for *ciPLV*. To gain a better understanding of the performance of different methods, we estimated the true positive and false positive connections captured by applying different aggregation methods. This analysis revealed that *Mean after* and *Max after* captured a larger number of true connections, but also a larger number of false positives. In other words, *Mean after* and *Max after* showed higher sensitivity for connectivity but lower specificity.

As for the AUC values, we found that there were differences between *PLV* and *ciPLV*. For *ciPLV*, *Mean after* and *Max after* performed better (with higher AUC) when only the center of the patch was considered, while for *PLV*, *Mean after* had the highest AUC regardless of the parcel size. While different strategies may be more or less appropriate depending on the specific scenario, and it may be difficult to determine which aggregation method performs best in real-world situations, our results suggest that *Mean after* and *Max after* may be preferable, however with the expense of longer computation time and higher rates of false positives. In addition, while PCA provided maps that were very different from all other extraction methods in real data, simulated data showed similar behaviour to *Mean before*. Overall, we believe it would be reasonable to choose *Mean before* when higher specificity and shorter computation time are desired or *Mean after* when greater sensitivity is desired.

The effects of extraction methods on connectivity analysis have been extensively studied in the field of fMRI. Previous research has shown that dimensionality reduction at the level of an ROI can result in the loss of important information (Basti et al., 2019). As a result, it has been suggested that multi-dimensional connectivity methods, also known as multivariate connectivity, may provide more accurate results [Geerligs et al., (2016); for a recent review see Basti et al., (2020)]. Applying dimensionality reduction strategies at the seed level to obtain a single time-course is recommended only in the case of homogeneous ROIs, which is rare in M/EEG due to cortical folding that often results in opposite directions for reconstructed source space time-courses. Several advanced multivariate approaches that integrate the computation of connectivity and the extraction of unique patterns of relationship among brain regions have been developed (Basti et al., 2018; Ewald et al., 2012). A recent MEG simulation study compared standard procedures based on the identification of a representative time-course or ROI (average, centroid, largest power, first PCA component, and first Kosambi-Hilbert torsion component) against a multivariate approach and concluded that the latter is remarkably better (Bruña & Pereda, 2021). Another study compared three approaches to extract the time-course of the ROI (centroid, first PCA component, average) and two multivariate approaches, namely the average of the pairwise connectivities across all elements of the ROIs (corresponding to the *Mean after* of our study) and the root-mean-square (RMS) of all pairwise connectivity in an EEG-MEG study (Bruña et al., 2023). The results showed that multivariate procedures (*Mean after* and RMS post) performed better based on the concordance between EEG and MEG source-based connectivity. However since no ground truth was available in that study, it cannot be ruled out that some of the results are driven by the inherent differences of EEG and MEG. Despite the development of new multivariate approaches, many M/EEG researchers continue to rely on default approaches built in among most commonly used toolboxes for connectivity analyses (i.e., Brainstorm, Fieldtrip, MNE python) (Gramfort et al., 2014; Oostenveld et al., 2011; Tadel et al., 2011). Within these toolboxes, the most used extraction strategy is the average time course of the ROI. The use of PCA for dimensionality reduction has received conflicting evaluations. Some studies suggest that PCA may work better than averaging the seed signal in cases when a process might be captured better by some voxels than others (Basti et al., 2020). Other studies suggest that PCA may be biased towards weighting more low frequencies at the expense of high frequencies (Chalas et al., 2022). A recent simulation study demonstrated that PCA appears to be the best technique, particularly when a fixed number of components is chosen across areas. However, this is only true when PCA is coupled with other optimal pieces of the pipeline, including specific inverse solutions and connectivity measures (Pellegrini et al., 2022). Additionally, another MEG study in clinical and control populations revealed that PCA is outperformed by the centroid of the ROI and that the choice of the aggregation procedure matters in healthy subjects as well as in patient populations (Dimitriadis et al., 2018).

## Limitations

This study has some limitations that should be acknowledged. First, the generalizability of the results may be limited as it was not feasible to compare all possible strategies available. Instead the study focused on a set of common and practical scenarios. Additionally, the effect of flip sign was not investigated for the procedures of aggregation prior to computing connectivity, even though this is a popular approach to limit the cancellation effect (Ahlfors, Han, Belliveau, et al., 2010; Lai et al., 2018). The influence of flip sign on the estimation of connectivity remains unclear and requires further systematic investigation.

Another limitation is that the study only focused on resting-state data. While many studies have shown that some patterns of connectivity remain consistent between rest and tasks, recent literature suggests that individual variations should be considered (Colenbier et al., 2023). Therefore, the generalizability of the results to task-based data may be limited. Finally, while the use of realistic simulations allowed for the assessment of different procedures, ideal analysis would be performed on invasive EEG data. This is planned future direction, in order to provide a more comprehensive evaluation of the different aggregation procedures.

## Conclusions

The current study highlights the critical importance of the aggregation procedures in determining accurate connectivity estimations, in both real and simulated data. The choice of the extraction method has a great impact on the connectivity output. Differences are higher for ciPLV than PLV, with a similar pattern across frequency bands. Further, larger is the ROI the higher is the difference across connectivity outputs obtained with different strategies. Overall to obtain higher accuracy (higher sensitivity at the expense of lower specificity), *after* aggregation procedures (*Mean after*) are recommended in connectivity analyses, even though they are computationally demanding. Given the significant differences across aggregation methods, it is essential to exercise caution when comparing studies that employ different methods.

## Open access resources

Code to replicate and use clustering analysis is available here: https://github.com/jrasero/connectivity-measures-clustering

MEG resting state data will be openly accessible on San Camillo Open access portal. The zipped file contains all the t-maps of the contrasts across extraction methods, ROIs, frequency bands. Maps relative to different connectivity measurements are grouped in folders. Each folder also contains the .gii file for the reference cortex surface. You may use different software to visualize these maps.

If you wish to visualize the maps with <u>Brainstorm</u>:

1. Go to Default Anatomy. Right Click -> Import surface. Select Group_analysis_cortex.gii.
2. Go to the Group_analysis data folder (you should create one if not present). Go to Common Files. Right Click -> File -> Import Source Maps. Select the t-maps to be visualized.

For any additional info, please get in touch with the corresponding author.

## Notes

https://github.com/jrasero/connectivity-measures-clustering

### Competing Interest Statement

The authors have declared no competing interest.

### Summary of Updates

Again we have updated the co-authors list, the content of the manuscript stayed the same.

