## supplementary material for "The impact of ROI extraction method for MEG connectivity estimation: practical recommendations for the study of resting state data"

### Table of Contents

***Participants demographics*** ***1***

***Aggregation procedures offered by the most common toolboxes for EEG/MEG data analysis.*** ***2***

### Participants demographics

Below is a description of the demographic information pf the recruited participants. Note that some subjects have had multiple eyes closed sessions.

Table S1. Demographic details of the recruited sample

| | **Subjects (N=84)** |
|---|---|
| **SESSIONS** | |
| Eyes closed | N=113 |
| **AGE** | |
| Mean (SD) | 29.0 (7.14) |
| Median [Min, Max] | 27.0 [21.0, 58.0] |
| **EDUCATION in years** | |
| Mean (SD) | 17.8 (3.17) |
| Median [Min, Max] | 18.0 [13.0, 26.0] |
| Missing | 15 (17.9%) |
| **HANDEDNESS** | |
| left | 4 (4.8%) |
| right | 80 (95.2%) |
| **SEX** | |
| female | 66 (78.6%) |
| male | 18 (21.4%) |

### Aggregation procedures offered by the most common toolboxes for EEG/MEG data analysis.

MNE Python offers the option to compute time course across labels or ROIs according to different modes: 1) maximum value across vertices at each time point ('max'); 2) average across vertices at each time point within each label or ROI ('mean'); 3) average across vertices at each time point within each label or ROI after having flipped the time-courses with an opposite orientation to the dominant one in the parcel ('mean_flip'). The default behaviour is usually represented by the 'auto' option, that either uses 'mean' or 'mean_flip' to reduce signal cancellations. FieldTrip in ft_sourceparcellate has different options: 1) 'mean', 2) 'median', 3) 'eig' (largest eigenvector), 4) 'min', 5) 'max', 6) 'std' (standard deviation). Brainstorm offers the following options: 1) mean before, corresponding to the average across all time series of an ROI prior to computing connectivity; 2) PCA before, which takes the first mode of the PCA decomposition of all time series prior to the connectivity estimation; 3) Max after, computes connectivity for each time series of the ROI prior to selecting the maximal connectivity value; 4) Mean after, as Max after but takes the mean rather than the Max.

### Simulation false positive rates across the two connectivity measures for different path sizes

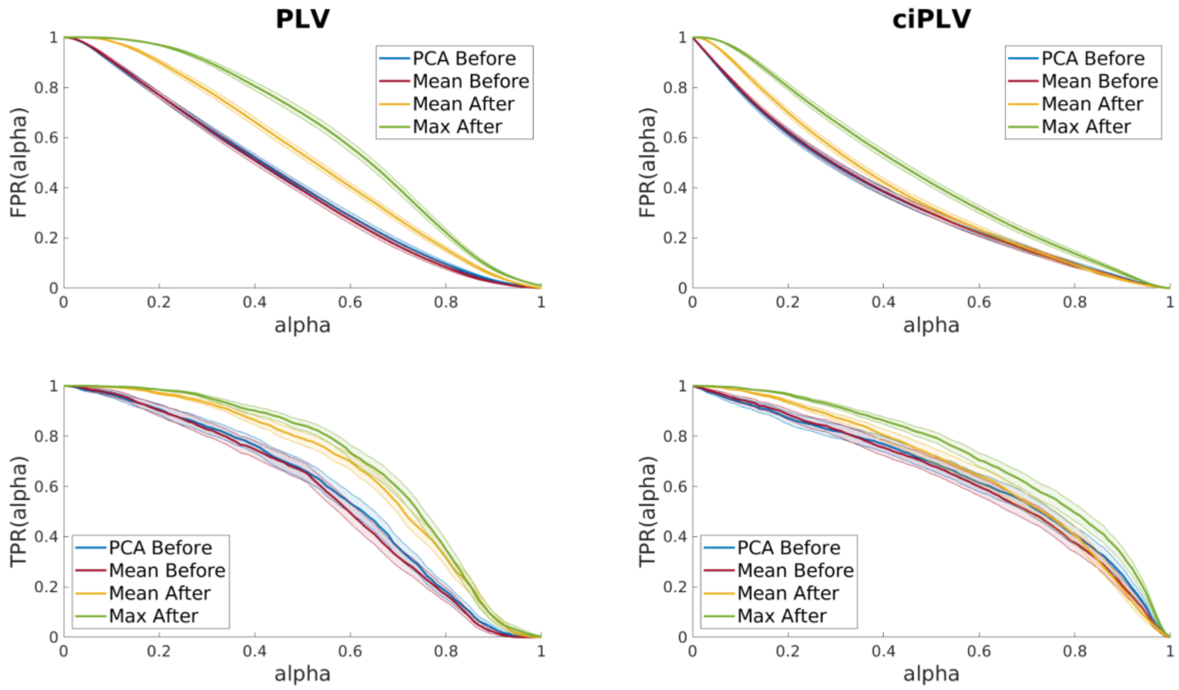


**Supplementary Figure S1.** False Positive Rates (*upper row*) and True Positive Rates (*lower row*) models of the two connectivity measures (PLV -left- and ciPLV -right-) extracted via the four different scout functions. Plots show mean and standard error of the mean across 68 simulated neural activity so that P1 contains the source-space points that are less than 2.5 cm apart from v1. Additionally, only points that belong to v1's AAL's (Desikan Killiany) ROI are retained.

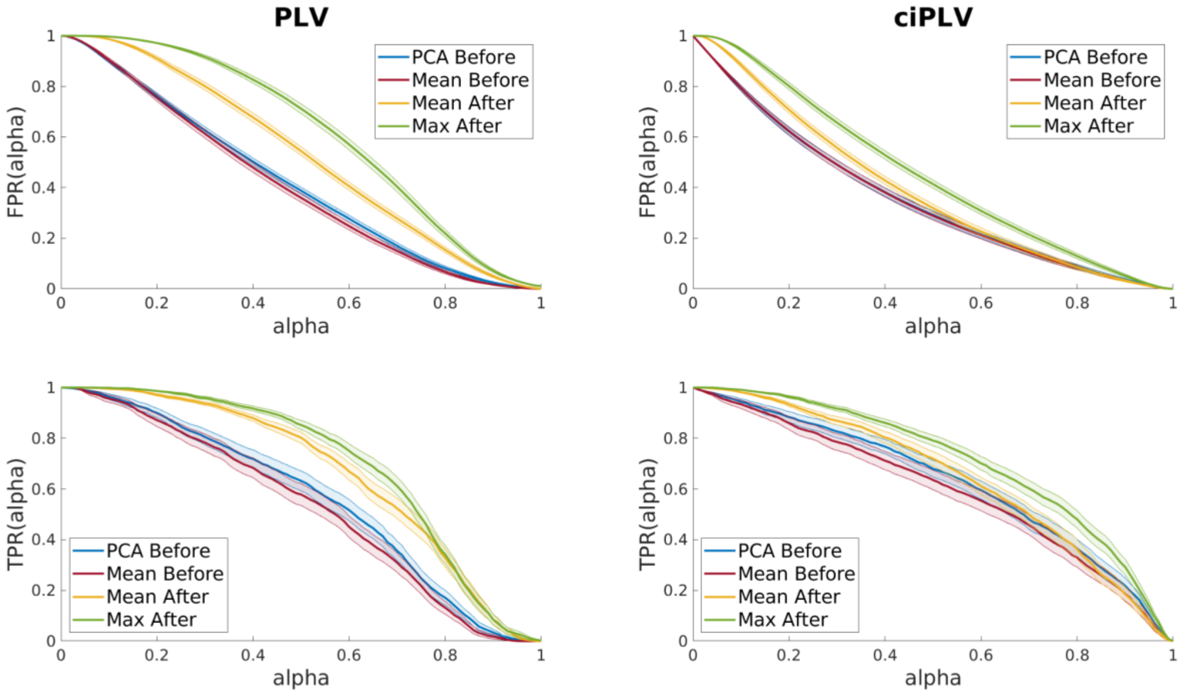


**Supplementary Figure S2.** Same as in Figure S1, but considering a patch around P1 which is half of the original size.
